# Developmental defects in ectodermal appendages caused by missense mutation in *edaradd* gene in the *nfr* mangrove killifish, *Kryptolebias marmoratus*

**DOI:** 10.1101/2024.04.24.591042

**Authors:** Hussein A. Saud, Paul A. O’Neill, Brian C. Ring, Tetsuhiro Kudoh

**Affiliations:** Department of Pathological Analyses, College of Science, University of Basrah, Basrah, IQ; Biosciences, College of Life and Environmental Sciences, University of Exeter, Exeter, EX4 4QD, UK; Department of Biology, College of Science & Math, Valdosta State University, 1500 N. Patterson St., Valdosta, GA 31698

**Keywords:** edaradd, eda, Kryptolebias marmoratus, Aphanius dispar

## Abstract

From an ENU-mutated mangrove killifish line R228, we have identified and isolated a novel mutant line, no-fin-ray/*nfr* in which fin ray development is largely reduced. Besides the reduction of the fin, the *nfr* mutant also exhibited other phenotype associated with ectodermal cell lineages including loss of scales, deformation in the gill structure such as decreasing the number of gill filaments, the reduction in the number of jaw teeth, pharyngeal teeth and gill rakers. Illumina RNAseq with 12 embryos each from mutants, siblings and the parental WT strain Hon9 identified a mutation in the *edaradd* in a highly conserved C-terminal death domain. Edaradd is known as a cytoplasmic accessory protein for the Ectodysplasin A (*EDA*) signalling pathway. To confirm the crucial role of *edaradd* during fin development, CRISPR RNAs were designed to knock out the gene in another killifish species, Arabian killifish. Indeed, Arabian killifish *edaradd* crispants showed a potent reduction of the fin development with 100% frequency. Furthermore, *EDA* crispants also showed identical phenotypes to that of *edaradd* crispants, confirming the fin defect in the mutants/crispants is caused by the signalling pathway of the *EDA* in the killifish species. These data demonstrate a powerful genetic approach using isogenic self-fertilising mangrove killifish as a tool for identifying mutants and their mutation, and revealed the crucial role of *edaradd* in the fish fin development and other ectoderm derived epithelial tissues.

## Introduction

The self-fertilising mangrove killifish, *Kryptolebias marmoratus*, is a powerful model for a variety of genetics studies. There are two sexual forms, hermaphrodite and male. Hermaphrodites can generate fertilised eggs by internal self-fertilisation and they also can sexually reproduce by outcrossing male fish (Mackiewicz *et al*., 2006). Laboratory strains were established by successive self-fertilisation for many generations (over 20 years) leading to the generation of highly isogenic and inbred lines (Tatarenkov *et al*. 2010; Moore *et al*., 2012). Self-fertilisation and isogenicity makes this animal model extremely unique and useful for generating mutant lines and identifying the mutated gene(s) (Sucar *et al*. 2016; Saud *et al*., 2021). Firstly, when a mutated F1 fish contains a mutation in the genome, as this fish is a self-fertilising hermaphrodite possessing ovary and testis in the same body, the fish already produces eggs and sperms heterozygous for the same mutation. These fish are able to generate F2 homozygous mutant embryos within predictable Mendelian ratios (Sucar *et al*. 2016). Therefore, it is not necessary to separate groups of siblings in individual tanks and screen for carriers of the same mutation from two both sexes. Secondly, due to the isogenicity of the laboratory lines, genetic variations between two complementary chromosomes are highly limited, facilitating instant identification of mutated genes by a single run of sequencing with a small number of mutant, siblings and parental wild type strains, avoiding the need for genetic mapping (Saud *et al*., 2021).

Ectodermal dysplasia (ED) is a hereditary disease having defects in development of hair, teeth, secretory glands and digits (Kowalczyk-Quintas and Schneider, 2014). From family genetic analyses, it has been discovered that one of the ED subtypes, hypohydrotic ectodermal dysplasia (HED) in patients that have a mutation in one of the three genes, ectodysplasin A/*eda, edar* or *edaradd* (Chassaing *et al*., 2010). HED is diagnosed through chronic diseases including hypotrichosis, hypodontia and anhidrosis resulting in the reduction or absence of eccrine sweat glands, hair follicles and teeth, and defective formation of salivary, mammary and craniofacial glands (Chassaing *et al*., 2010).

*Eda* encodes a TGF-beta super family signalling molecule. *EDA* protein binds to the receptor, *EDAR*, and in turn *EDAR* is associated with cytoplasmic protein *EDARADD* (Sadier *et al*., 2015). *Edaradd* encodes a protein that is considered as adaptor interacting with EDAR. EDAR/EDARADD complex transduce the signal via TAB2, TRAF6 and TAK1 to activate transcription factor NF-kappaB to target gene expression involved in ectodermal structural development (Morlon *et al*., 2005).

Similar to human patients, in the mouse, *edar*/*edaradd* mutations cause the disappearance of hair follicle and eccrine sweat gland and disorder in the glands of salivary and mammary craniofacial glands (Del-Pozo *et al*., 2019 and Kuramoto *et al*., 2011). Furthermore HED mutant mice are affected by persistent rhinitis and otitis that arise from disorder in the mucociliary role (Azar *et al*., 2016).

In fish species, *Eda* and *edar* gene functions have been investigated in zebrafish and medaka. In zebrafish, *eda* and *edar* mutations (*nackt* and *finless* respectively) were identified by Harris *et al*. (2008) causing the disappearance of fins and scales and reduction in the number of pharyngeal teeth. Iida *et al*. (2014) also reported a nonsense mutation in *eda* in m*eda*ka fish called *alf* presenting coiled caudal peduncle without fin rays missing in all median and paired fins with a deformed skull, scales and teeth.

Though the crucial role of *eda* and *edar* have been studieed in these two model fish species, the role of *edaradd* was not fully known in the teleost fish. There is only one report of *edaradd* morpholino gene knock down study showing reduced development of the jaw and teeth (Sadier *et al*., 2015). However the phenotype in the fin development was not reported in the loss of function of *edaradd* in fish species thus far. As morpholinos can be degraded after a few days of application it is conceivable that the phenotype in the fin development may not be fully affected by morpholino experiments. Therefore loss of function studies involving genetic mutations of *edaradd* in fish models have been needed.

The first ENU mutagenesis applied to the self-fertilising mangrove killifish generated a variety of mutant phenotypes (Moore *et al*., 2012, Sucar *et al*., 2016). Among these, two lines R109 and R228 were sequenced and identified the mutations in the genes, *noto* and *msgn1* causing the phenotype of the shorttail/R109 and balltail/R228 respectively (Saud *et al*., 2021). While maintaining the mutant lines, we identified another mutant phenotype that we named no-fin-ray (*nfr*) that was found from the R228 progenitor lineage. Here we report the *nfr* has a mutation in the *edaradd* gene and we show here a crucial role in the fin development and other ectoderm derived tissues including teeth, jaw and scale.

## Materials and methods

### Maintenance of the mangrove killifish and Arabian killifish

The mangrove killifish, R228 strain and parental wild type strain, Hon9 were maintained under the same laboratory conditions with 15ppt artificial sea water and aeration at 26° C ±1, 12 h light: 12h night photoperiod. Fish were fed on live *Artemia* once a day. Each fish were individually separated in 1.5L containers. Water was changed once weekly. R228/*nfr* strain was provided from Ring’s lab (Sucar *et al*., 2016). Eggs were collected by natural spawning from mature fish in both WT, mutant, and heterozygous carrier siblingstrains.

The Arabian killifish wild type strain was maintained with 35ppt artificial sea water in 40L tanks as a small group with circulation and aeration at 26° C ±1, 12 h light: 12h night photoperiod. Egg chambers were placed at the bottom of the tank before the day of egg collection and eggs were collected by natural spawning in the morning 30min to 1 hour after the start of the light period.

### RNA extraction and analysis

Total RNAs were extracted used Qiagen RNA extraction kit from twelve embryos from the mutants, siblings and from wildtype (Hon9 strain) for the stages 16-18. Quality of RNAs were tested by Agilent RNA 6000 Nano Kit then sequenced using an Illumina HiSeq 2500 v3 next generation sequencer. Sequencing was conducted with 100 bp paired end reads on one lane. Sequencing adaptors firstly was trimmed then the low quality flanks(<Q20) and short segments were removed using fastq-mcf v1.1.2-537. Assembly of transcriptomes were reconstructed by Trinity v2.2.0 (Haas *et al*., 2013). Variants were identified using KisSplice v2.4.0-p1 (Lopez-Maestre *et al*., 2016) into k-mer size of 53, then mapped to de novo transcriptomes by BLASTn v2.5.0 (Altschul *et al*., 1990). Transcripts were annotated using blast-NCBI, and to select the best hits, threshold of 1e−4 value to identify the suspected candidate genes. According to Saud *et al*., (2021), 100% of reads for mutants sample compared with 0% of reads for the wildtype samples, greatly simplifies mutantation identification.

### Skeletal staining with Alcian blue and Alizarin red

Samples was fixed with PFA, then washed with deionized water (2 times), and several times with DI water (2 hr). Next water was replaced with Alcian blue (2 hr) and placed into a series of ethanol washes (75%,50%,25%) each for ½ hr, and washed with 30% of tetra borate (2 times, 5 min. each.Next, the samples were transferred to Trypsin/sodium borate (0.12/30%) overnight, then washed with 2% KOH (2 times, 5 min. each) and replaced with Alizarin red (0.002% in 2% KOH) for one day. To bleach, embryos were placed into bleaching mixtures (3 of 0.5% KOH + 1 Glycerol + 10 µl H2O2) 3-4 hr, and for clearance, the embryos were treated with a series of 0.5% KOH: Glycerol (1:1, 1:3, 100% Glycerol,2 hr each). Finally specimens were stored in 100% Glycerol with a few thymol drops to prevent decay (Dingerkus and Uhler, 1977).

### Gene Knock out of *edaradd* and *eda* in the Arabian killifish

Two CRISPR RNAs were designed from *edaradd* (AAGAACTTTGCAAGCCGTTG and CTGCTAGAGCACCGGACCCA) and *eda* (ACCAACCACACGACCTTCCT and TTCAACACCTGCTACACGGC). Two crRNAs (final 50ng/ul each, Integrated DNA Technologies; IDT) and tracr RNA (final 100ng/ul, IDT), Cas9 nuclease protein (final 1ug/ul, IDT), 0.25% phenol red (Sigma) were dissolved in dilution buffer (IDT). Approximately 1 nl of the above crRNA mixture were injected into 1-cell stage Arabian killifish eggs. Embryos were cultured at 28’C for 11 days (the day of hatching)and wereanaesthetised with tricaine and imaged using a Nikon SMZ1500 stereo microscope.

## Results

### Paired and single fins were deleted in *nfr* mangrove killifish

*Nfr* mutant presented severe phenotype showing deletion in all fins (Fig.1). At early stages of development (specifically at stages 28 to 29) fin formation starts to emerge (Mourabit *et al*., 2011).

**Figure 1.**
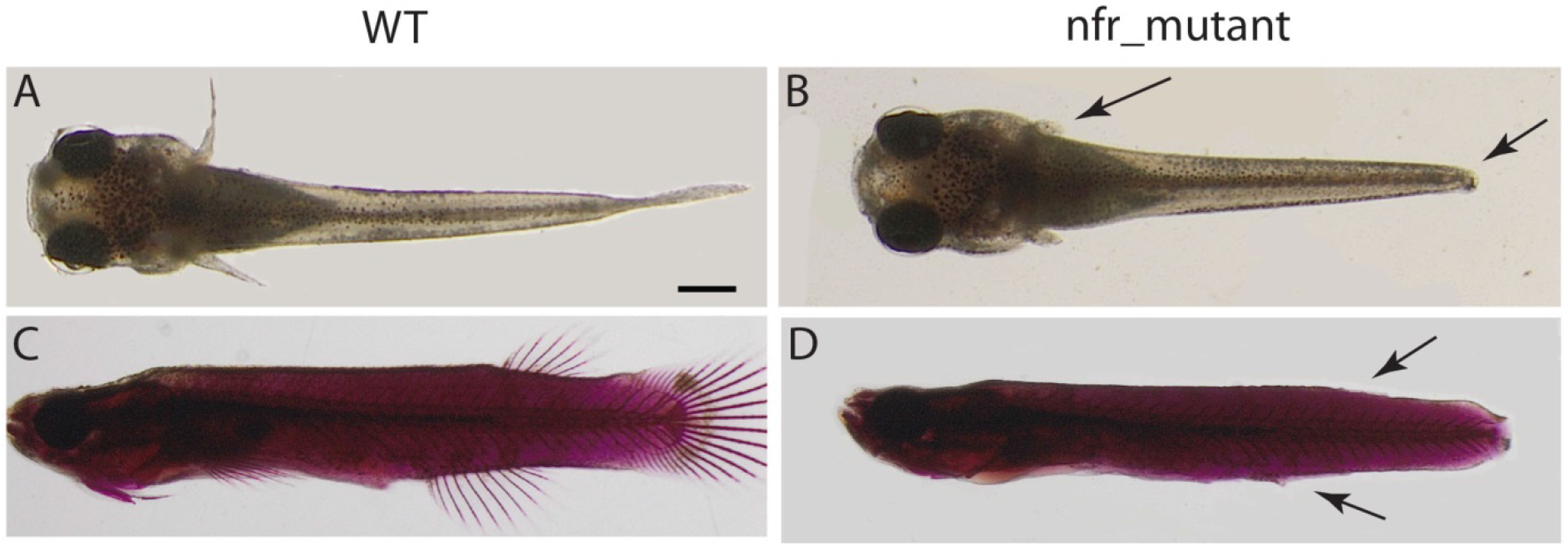
The mangrove killifish *nfr* mutant shows substantial reduction of the fin development in the post hatching larvae and juvenile fish. A and C are wildtype, B and D *nfr* mutants. A and B are post-hatched larvae and C and D are juvenile fish. Arrows refers to missing caudal and pectoral fins (B), dorsal and ventral fins (D). Both C and D are stained with alizarin red and alcian blue.

### Scales missing and mouth parts deformation in *nfr* mutation

The phenotype examination in the skin and mouth parts revealed skin free of scales and decreasing in the number of teeth in jaws and pharyngeal teeth (Fig.2). It was observed remain the big teeth with missing the small one in both jaws and pharyngeal teeth. In addition, there was an obvious and dramatic decrease in the number of gill filaments and gill rakers.

**Figure 2.**
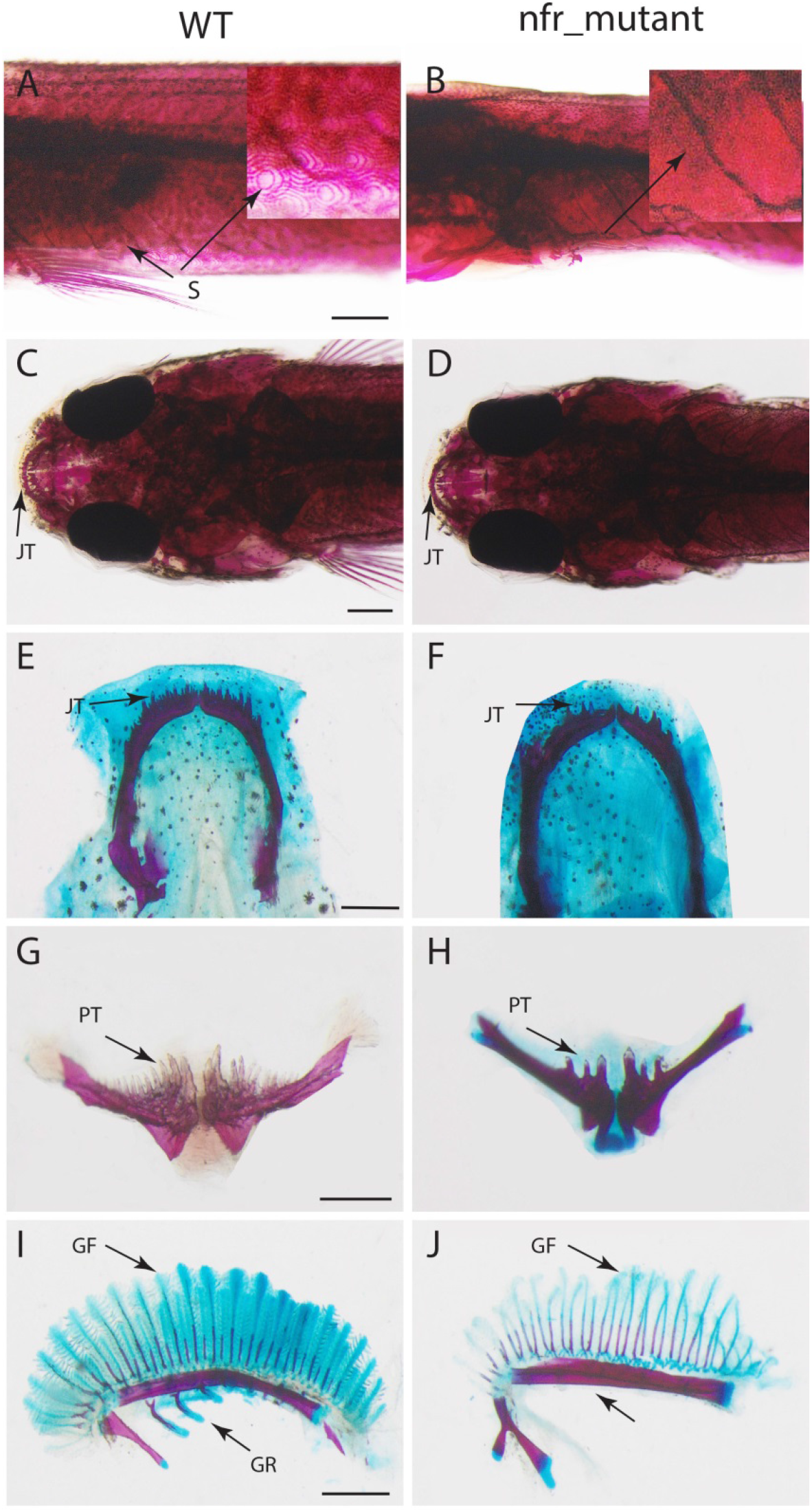
Scales, teeth and gills in mangrove killifish. A,C,E,G and I wildtype; B,D,F,H and J *nfr* mutant. S: scales (A), no scales in B; JT: jaws teeth, in D and F number of teeth were reduced compared with wildtype (C and E); PT: pharyngeal teeth are reduced in H compared with wildtype (G); number of gill rackers (GR) and gill filaments (GF) are dramatically decreased. All parts stained with alizarin red and alcian blue.

### Nfr is missense mutation of edaradd

To identify the *nfr* mutation, *nfr*, their sibling and WT embryos were analysed with RNAseq. Four duplicates of three embryos/group from each category were sequenced with one lane of Illumina HiSeq 2500 100bp paired end read. From the variations identified, the one which is 100% enriched in the mutant group, 0% in the WT group and intermediate number in the sibling group were searched, leading to the identification of 19 potential sequence variations. Among these, only three variants were in the protein coding region: One was synonymous variation and other two were non-synonymous. These two non-synonymous mutations were seen in the genes, *edaradd* and carboxypeptidase-O like gene. In the sibling group, if the variation is responsible for the phenotype, the WT sequence is expected to be twice abundant than mutant path. Carboxypeptidase-O like gene showed far larger read of mutant sequence implied that this was not responsible for the *nfr* mutant phenotype. Consequently candidates for the *nfr* mutation were narrowed down to the single gene, *edaradd* (Fig.3). In the *nfr* mutant, Arg 192 is substituted to Cysteine. In the *nfr* mutant, Arginine 192 is within the Death Domain and is a highly conserved amino acid among orthologs in vertebrate species (Figure 3). A missense mutation in a highly conserved amino acid in the functional domain suggested that the mutation would cause a severe loss of function of the protein.

**Figure 3.**
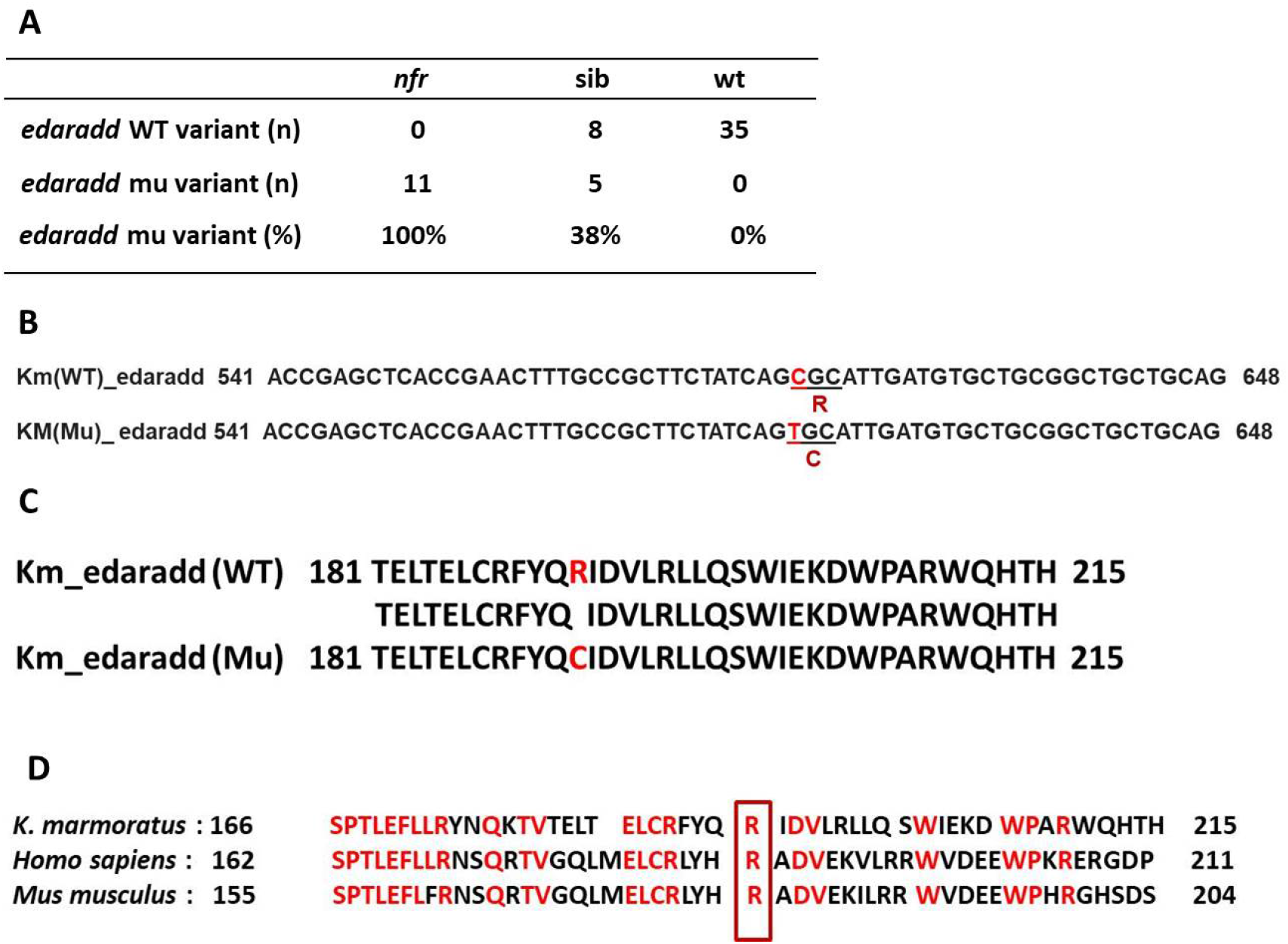
A: The mutation in the *edaradd* was 100% enriched in the *nfr* mutant embryos, 38% in the siblings and 0% in the WT embryos.. B: Point mutation at loci 574 in nucleic acid base cytosine changed it to thymine in Km-*nfr* mutant. C: The resulting missense mutation changes the amino acid Arginine to Cysteine in the mutant. D: The mutation occurs in the highly conserved Arginine that is in the C-terminal death domain common to both the mangrove killifish (Km), thehuman and mouse.

### Knock out *edaradd* in Arabian killifish phenocopies *nfr*

To confirm that loss of function of *edaradd* can cause the no fin ray phenotype, two CRISPR RNAs were designed from the *edaradd* gene and administered to the Arabian killifish model. Due to internal self-fetilization prevalent in the mangrove killifish it is difficult to generate sufficient number of 1-cell stage embryos for crisper injections from a limited number of adult fish. So we used the Arabian killifish to recapitulate the nfr phenotype. The CRISPR/Cas9 genome gene knockout treatment was highly effective in the Arabian killifish. Indeed the crispants of *edaradd* showed a clear loss of caudal fin 11dpf embryos/larvae (the hatching stage) with 100% frequency (Fig. 4). At this stage, defects in the pectoral fin were not that obvious.

**Figure 4.**
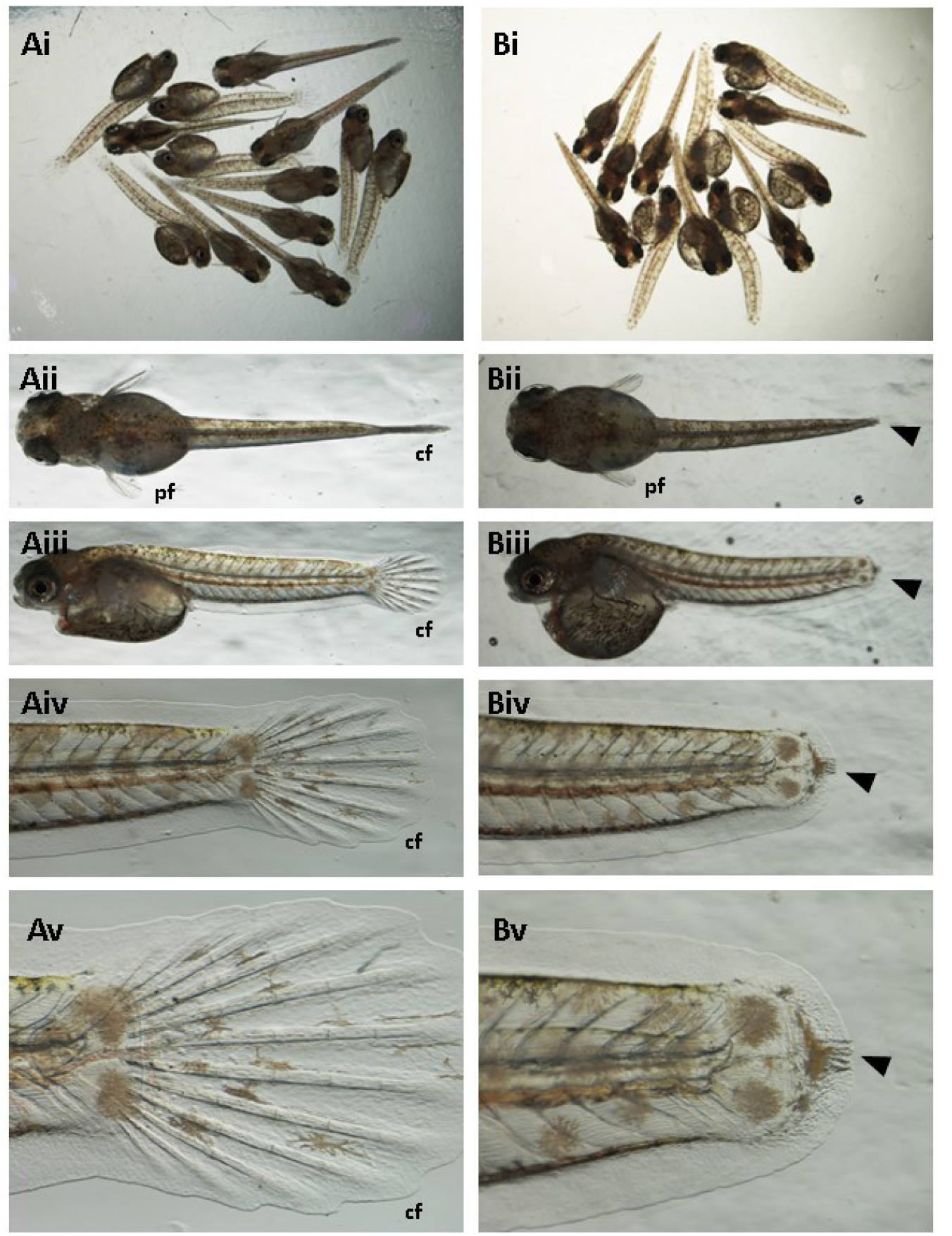
*Edaradd* crispants show loss of the caudal fin with 100% frequency. Mixture of two CRISPR RNAs for *edaradd* was injected into the 1-cell stage Arabian killifish eggs. Phenotypes were analysed at 11dpf (hatching stage) showing loss of the caudal fin (cf) in all crispant embryos (Bi to Bv) as compared to WT controls (Ai to Av). At this stage, pectoral fin (pf) development was not significantly affected.

### *Edaradd* loss of function shows similarity to the ones for *eda*

It has been known that *edaradd* act as the downstream cytoplasmic adaptor molecule of the *EDA*/*EDAR* signalling pathway. To confirm that *edaradd* is indeed acting along this pathway in the teleost species during fin development, the upstream gene, *eda*, was also knocked out by injecting CRISPR RNAs in the Arabian killfish. Although the percentage of the embryos showing abnormalities were lower in the *eda* crispants (70 to 80%, Fig. 5) as compared to the *edaradd* crispant treatment group (100%), we observed an indistinguishable morphological change ofcaudal fin loss at the hatching stage in both treatments (Fig. 4 & 5). Not only the loss of the caudal fin, but also the morphological details such as abnormal accumulation of pigment cells at the growing edge of the caudal fin was equally observed in the crispants of the two genes. As seen in the *edaradd* KO, *eda* KO also did not show a significant reduction of the pectoral fin at the hatching stage.

**Figure 5.**
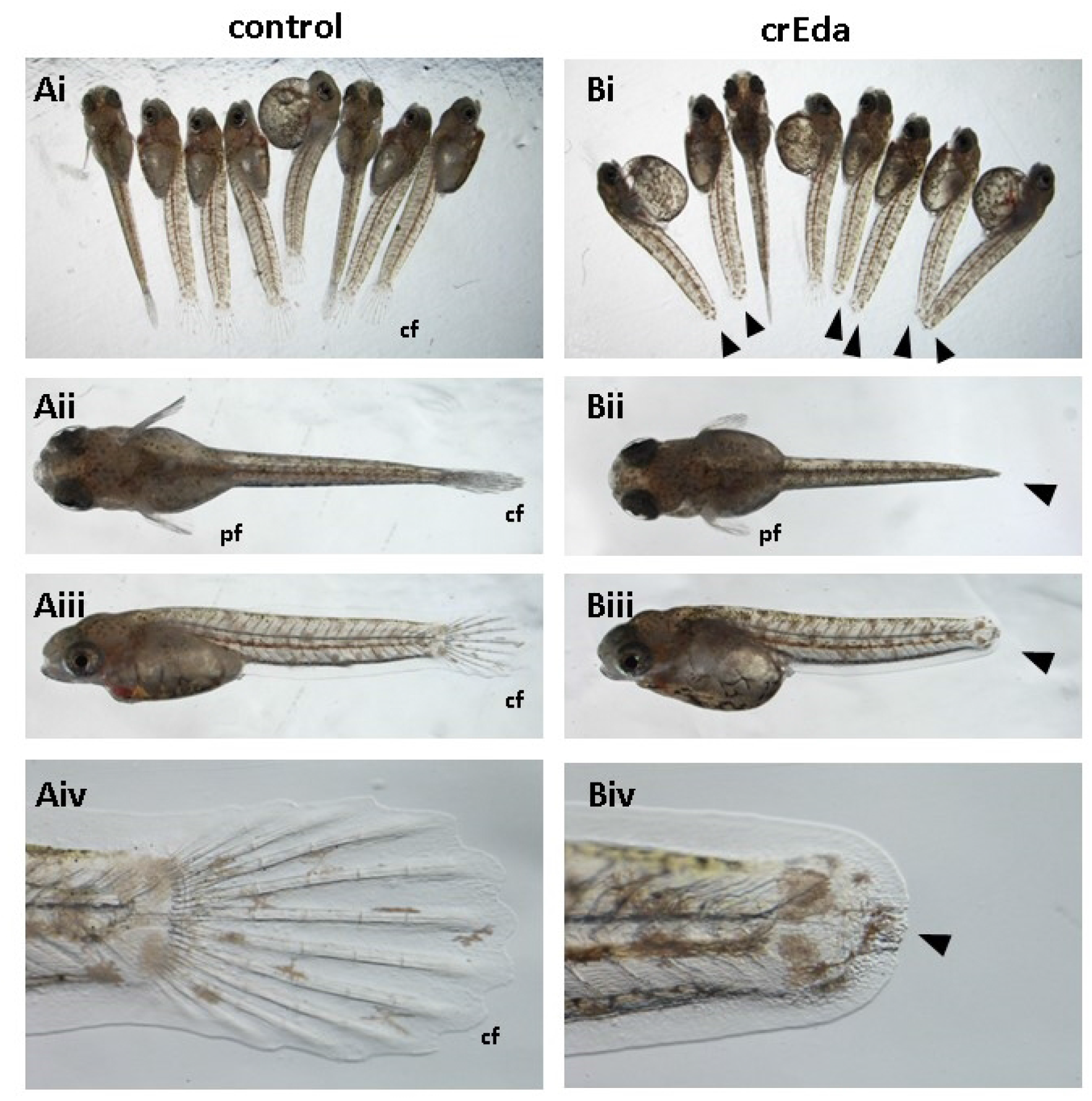
*Eda* crispants show loss of the caudal fin with 70-80% frequency (Bi to Biv) versus WT controls (Ai to Aiv). A mixture of two CRISPR RNAs for *edar* was injected into the 1-cell stage Arabian killifish eggs. Phenotype were analysed at 11dpf (hatching stage) showing loss of the caudal fin (cf) in the majority of crispant treated embryos. At this stage, pectoral fin (pf) development was not significantly affected.

## Discussion

Here we report identification of no fin ray mutant from a previous ENU mutagenesis screen utilizing the self-fertilizing mangrove killifish. The homozygous *nfr* fish are viable and can spawn eggs as normal with self-fertilisation with 100% frequency of *nfr* phenotype in the offspring forming a true breeding clonal mutant line. RNAseq analysis of the *nfr*, sibling and Hon9 (parental WT strain) with a small number of mixed embryos (4 duplicates samples) was sufficient to narrow down all variations to a single non-synonymous mutation that occurred in the *edaradd* encoding Arginine 192. The resulting missense mutation changed Arginine to Cysteine in the C-terminal death domain. Using the same method employed previously in the characterization and identification of two other mutants, *shorttail* and *balltail* as the mutations of *noto* and *msgn1* respectively (Saud *et al*., 2021), this method is a powerful tool for rapid identification of mutant alleles identified through forward mutagenesis. In all cases, this process of identifying corresponding mutations was highly simplified and straight forward for detecting single mutations in candidate genes. This isstrong evidence supporting the use of isogenic mangrove killifish strains as extremely useful for identifying mutations as the number of polymorphic variations are highly reduced. Even in the protein non-coding regions, we only identified 16 variants, suggesting that more subtle mutation phenotypes that are caused by mutations in the non-coding regions or promoter/enhancer regions may also be identified and characterised in future forward genetic screens.

This is the first report showing that an *edaradd* mutation causes the loss of fin phenotype in teleost fish species and we have confirmed it in two species, the mangrove killifish and Arabian killifish. The phenotype that was observed in this work with *edaradd* mutations was highly similar to the one reported in zebrafish with *eda* and *edar* (Harris *et al*., 2008) and m*eda*ka with *eda* (Iida *et al*., 2014). These findings indicate the pathway is widely conserved in teleost fish species. And in a broader context, the pathway is also conserved between fish and mammals (Pantalacci *et al*., 2008).

Appendages and limbs in fish represent the fins, but in human and other mammalians limbs form hands and feet.

*Nfr* in mangrove killifish resulted from a point mutation in the coding region at nucleotide 574 resulting in a missense mutation from cytosine to thiamine in the resulting protein. This particular missense mutation (allele) is a <u>nonsynonymous substitution</u> among others detected, and results in a change in a highly conserved amino acid not yet detected in other organisms.

To determine if the *nfr* allele functions similarly in another species, a crisper knock out of *edaradd* was performed in another genetic model species (Arabian killifish) belonging to the Cyprinodontidae family of Teleost fish. These experiments demonstrated that a similar phenocopy of *nfr* mutant could be recapitulated in a reverse genetic manor. Our results resulted in the disappearance of caudal, dorsal and ventral fins whereas the pectoral still developed, albeit with an abnormal shape. This experiment confirms the role of genetic pathways involved in fin development with slightly difference in defection and this is might be related to the presence of other genes (yet to be determined) that contribute to the development of appendages among vertebrate species. This is supported by the existence of some remnant rays in base of caudal fin.

Of further interest is the observation that caudal fin development was suppressed with loss of the fin ray and failure of pigment cell migration despite the size of pectoral fin was not significantly affected at this stage (hatching stage). It should be noted that in the caudal fin, although fin ray was not developed, the epidermal layer of the caudal fin fold seems fully developed (Fig.4). This may suggest that the role of the *eda*/*edar*/*edaradd* pathway is not directly involved in the fin epidermal development but upstream. In addition, the pigment cells (fluoroleucophore, Hashimoto *et al*, 2021) are accumulated at the tip of the tail inside of the caudal fin fold suggesting that neural crest cell differentiation to pigment cells in the caudal fin was not suppressed either. Considering this evidence, although all cell linage are missing in the caudal fin ray, it is possible that the role of this pathway is highly specific to a cell lineage such as cartilage cell differentiation as previously suggested and defects in other cell lineage may be a secondary consequence of the defect (Iida *et al*., 2014; Sadier *et al*., 2014; Lefebvre and Mikkola, 2014). Considering that early stage of fin development, such as fin bud formation is not affected and epidermal fin fold development is not affected neither, this may explain that the size of the pectoral fin at the hatching stage is not affected by the *edaradd* knockout.

Here we report a homozygous viable mutant strain in the mangrove killifish and identification of the mutated gene. In many cases, it is conceivable that homozygous viable mutants are not easy to maintain because even though they are viable, active mating behaviour might be compromised. However, by using a self-fertilising animal, it is possible to avoid the step of mating and lines can be established with relative ease. We have demonstrated this in subsequent genetic screens and more recently other researchers have added to repertoire of genetic tools in the mangrove killifish (Li *et al*. 2023). It is also possible to analyse subtle phenotypes (e.g. learning difficulty, aggression, altered facial structure, microcephaly) using this self-fertilising model. Geneticscreening of such weak phenotype at adult stage is easier as individual variation can be screened among established clonal lines. Therefore this model is poised for future studies of natural, mutagen induced, and CRISPR mediated genomic variants.

